# PETIL: Predicting Expansion of Tumor Infiltrating Lymphocytes for the Adoptive Cell Immunotherapy in Bladder Cancers

**DOI:** 10.1101/2025.10.15.682695

**Authors:** Kayode D. Olumoyin, Ahmet Murat Aydin, Sarah Bazargan, Brittany Bunch, Ibrahim Chamseddine, Aleksandra Karolak, Matthew Beatty, Shari Pilon-Thomas, Michael A. Poch, Katarzyna A. Rejniak

## Abstract

Adoptive cell therapy (ACT) with tumor-infiltrating lymphocytes (TIL) is a form of personalized immunotherapy that requires ex vivo expansion of autologous TILs and their reinfusion back into the patient. Predicting TIL expansion at the time of diagnosis may improve selection of patients that can benefit from ACT-TIL. It can also prevent high treatment-related costs and delays in treatment of patients whose cancer specimens would not yield successful TIL growth. We developed PETIL, a machine-learning model optimized for data of a medium size to determine a minimal combination of features (demographic, clinical, and biological specimen-based) that is predictive of expansion of TILs from a resected bladder cancer. We used a retrospectively identified set of data from bladder cancer patients at Moffitt Cancer Center for the training and testing cohorts. Additionally, we used data from a recent feasibility clinical trial at Moffitt Cancer Center as a blinded validation cohort. PETIL uses random forest method to identify a combination of robust predictive features, support vector machine model to determine the optimal classification hyperparameters, and Matthews correlation coefficient method to adjust the decision-boundary threshold for imbalanced data. Our model yielded AUC=0.740 for the testing cohort and AUC=0.857 for blinded validation cohort. Thus, our PETIL model optimized for data of medium size has favorable performance metrics for predicting TIL expansion from a given tumor.

**Authors Summary:** Treatment with autologous tumor-infiltrating lymphocytes (TIL) that are expanded ex vivo from a given tumor and then reinfused into the patient is a promising personalized immunotherapy. However, the TIL expansion takes about 4-6 weeks, thus developing tools that predict whether TIL growth will be successful can help to avoid delays in treatment of patients whose cancer specimens would not yield successful TIL expansion. Our Predictor of Expansion of TIL (PETIL) is a machine-learning model that uses patients’ demographic information, clinical tumor classification, and biological tumor specimen-based measurements to determine a minimal set of these data features that are predictive of TIL expansion outcome. We applied this model to data from bladder cancer patients collected at Moffitt Cancer Center and showed that PETIL has favorable performance metrics for the dataset of a moderate size. This computational predictor can support clinicians in determining which patients are candidates for TIL immunotherapy. The developed PETIL pipeline can also be adjusted to data from other solid tumors.

## Introduction

Bladder cancer is the fourth most common cancer among men and a leading cause of cancer death among men and women. The American Cancer Society estimates approximately 84,530 new cases of bladder cancer and 17,870 bladder cancer-related deaths in the United States in 2026 (1). Despite current therapies, 50% of patients with intermediate and high-risk localized non-muscle invasive bladder cancer fail bladder-sparing treatment. Additionally, recurrent or locally advanced tumors failing bladder-sparing treatment have an even worse prognosis, often requiring radical cystectomy, which is associated with several co-morbidity and changes in quality-of-life afterwards (2, 3). Further research using novel approaches is needed to treat this disease at every stage.

One major advance for treating solid tumors is the success of adoptive cell therapy (ACT) during which autologous tumor-infiltrating lymphocytes (TILs) are expanded and activated ex vivo and then reinfused into the cancer patient (4). Indeed, ACT with TIL has emerged as one of the most powerful therapies for unresectable metastatic melanoma and cervical cancer (5–7). Because bladder tumors have a high mutational burden corresponding to an increased number of neoantigens (8, 9), these cancer cells could be recognized by activated T cells at the tumor site, making the bladder cancer a potential candidate for ACT-TIL treatment. Prior studies by us and others showed that TILs expanded from bladder cancer recognize autologous tumor (10), and that expanding TILs ex vivo from the resected bladder tumors is feasible (11, 12). Moreover, bladder cancer provides a unique opportunity to deliver TIL intravesically by administering T cells through a catheter into the bladder directly to tumors. An active phase I feasibility clinical trial of intravesical adoptive cell therapy with TIL for high-grade non-muscle invasive bladder cancer (NCT05768347, Moffitt Cancer Center (13)) is showing that this treatment is well-tolerated.

One of the main steps in the ACT-TIL approach is the ability to expand tumor-reactive TIL to large amounts for reinfusion into patients’ bladders. Our previous work (12) showed that about 70% of collected primary bladder tumors were capable of TIL expansion. However, this process takes several weeks. Therefore, it would be beneficial to prognosticate early whether a patient will benefit from TIL therapy to allow for better patient stratification. One approach is using quantitative methods, such as machine-learning (ML) algorithms, to predict whether TILs can be expanded from resected tumors based on individual patient data. ML methods have been previously applied to several aspects of bladder cancer, such as evaluation of cancer stage in computed tomography urography (14); predictions of early recurrence of non-muscle invasive bladder cancer (NMIBC) based on histology images (15); and stratification of muscle-invasive bladder cancer (MIBC) patients into good or poor survival groups after chemotherapy (16). Here, we present the ML-based **P**redictor of the **E**xpansion of **T**umor **I**nfiltrating **L**ymphocytes (PETIL), a tool that can first learn from patient and tumor data collected in the clinic which data features are important for predicting TIL expansion, without the need to predefine which data categories to consider. Subsequently, this tool predicts a possible TIL expansion for individual patients (personalized predictions) allowing to determine whether ACT-TIL therapy could potentially treat an individual bladder cancer patient.

## Results

The goal of our studies was twofold. First, we aimed to identify a combination of data features among those previously collected in the clinic (and grouped as demographic, clinical, and specimen-based data) that were predictive of whether TIL can be expanded from that tumor. In this way, our model is optimized for local data, that is in contrast to other methods that use the predefined categories for data classification. Next, we used those data features to create and validate an ML-based predictor, PETIL, to stratify individual patients into *Yes-TIL* vs. *No-TIL* classes that define TIL expansion potential.

### Study design

The prospective database of 106 adult patients was created between 2015 and 2022 by collecting clinicopathologic and specimen information of bladder cancer patients undergoing surgery at Moffitt Cancer Center (Table 1). Data collection protocols were approved by the Advarra Institutional Review Board (MCC18142 and MCC20106). Informed consent was obtained from all patients prior to data collection. The initial screening identified the maximal subset of 60 patients with complete information on 15 commonly collected features that included (i) patients’ demographics: age at surgery, body-mass index (BMI), race, and smoker status; (ii) tumor clinical characteristics: clinical stage (cT), pathological stage (pT), pathological lymph node stage (pN), type of surgery, prior neoadjuvant chemotherapy (NAC), type of radical cystectomy histology (Histology), type of cystoscopic biopsy histology (Bx Histology), and cT/pT status; and (iii) specimen measurements: tumor sample weight, number of fragments plated for TIL expansion, and primary tumor digest count. The remaining 46 patients were missing data values in some of the considered features. Because individual features were not predictive of the TIL expansion status (Pearson correlation coefficients were between −0.21884 and 0.303202 for each of the 15 features, S1 Table), the combinations of features were considered, and the ML approach was applied. For every patient, information was recorded on whether the resected tumor sample caused positive TIL growth. Among the collected datasets, 68 tumors demonstrated TIL expansion (class *Yes-TIL*), and 38 did not (class *No-TIL*). This information was used for training and testing the PETIL predictor.

**Table 1.**
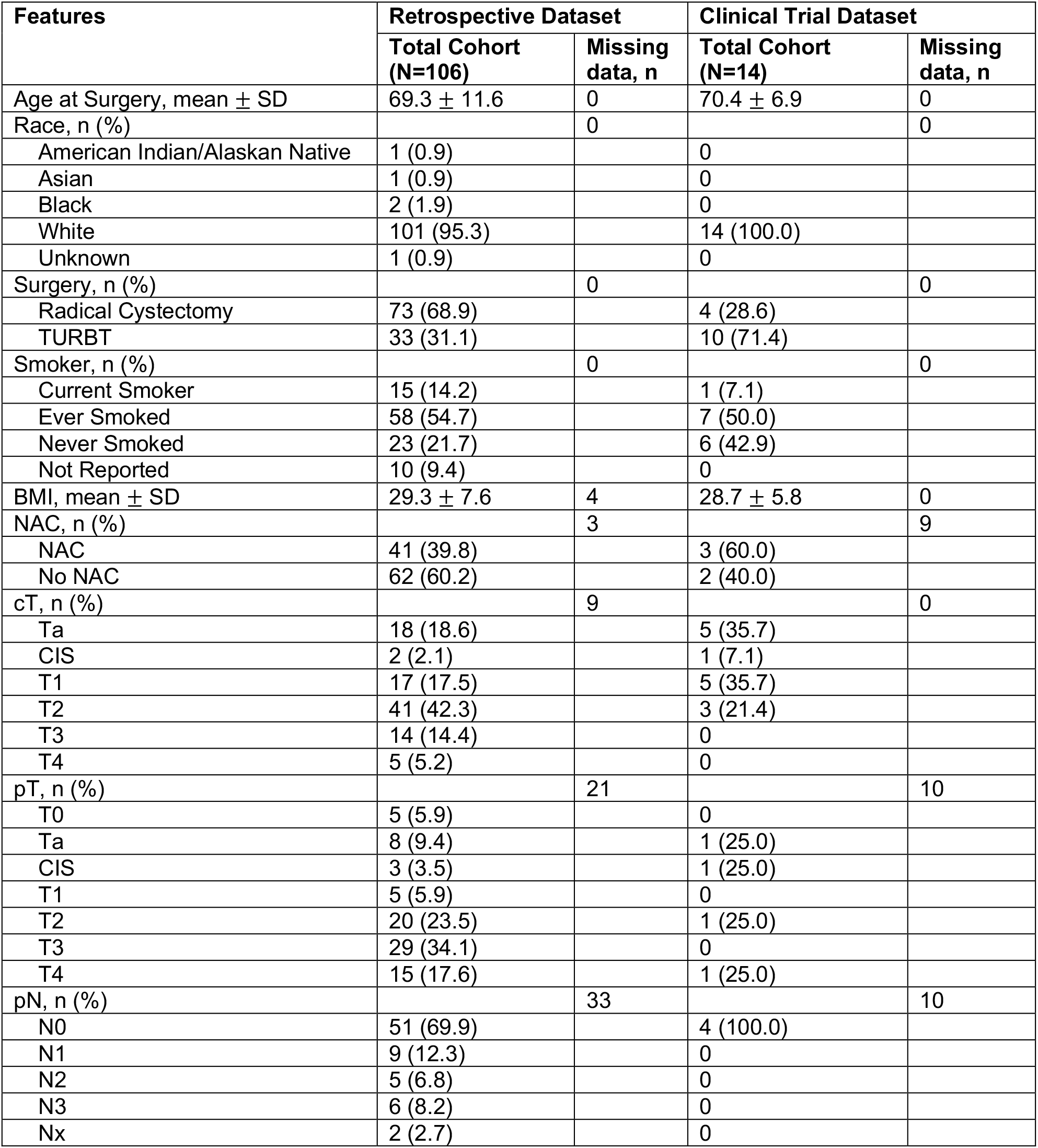
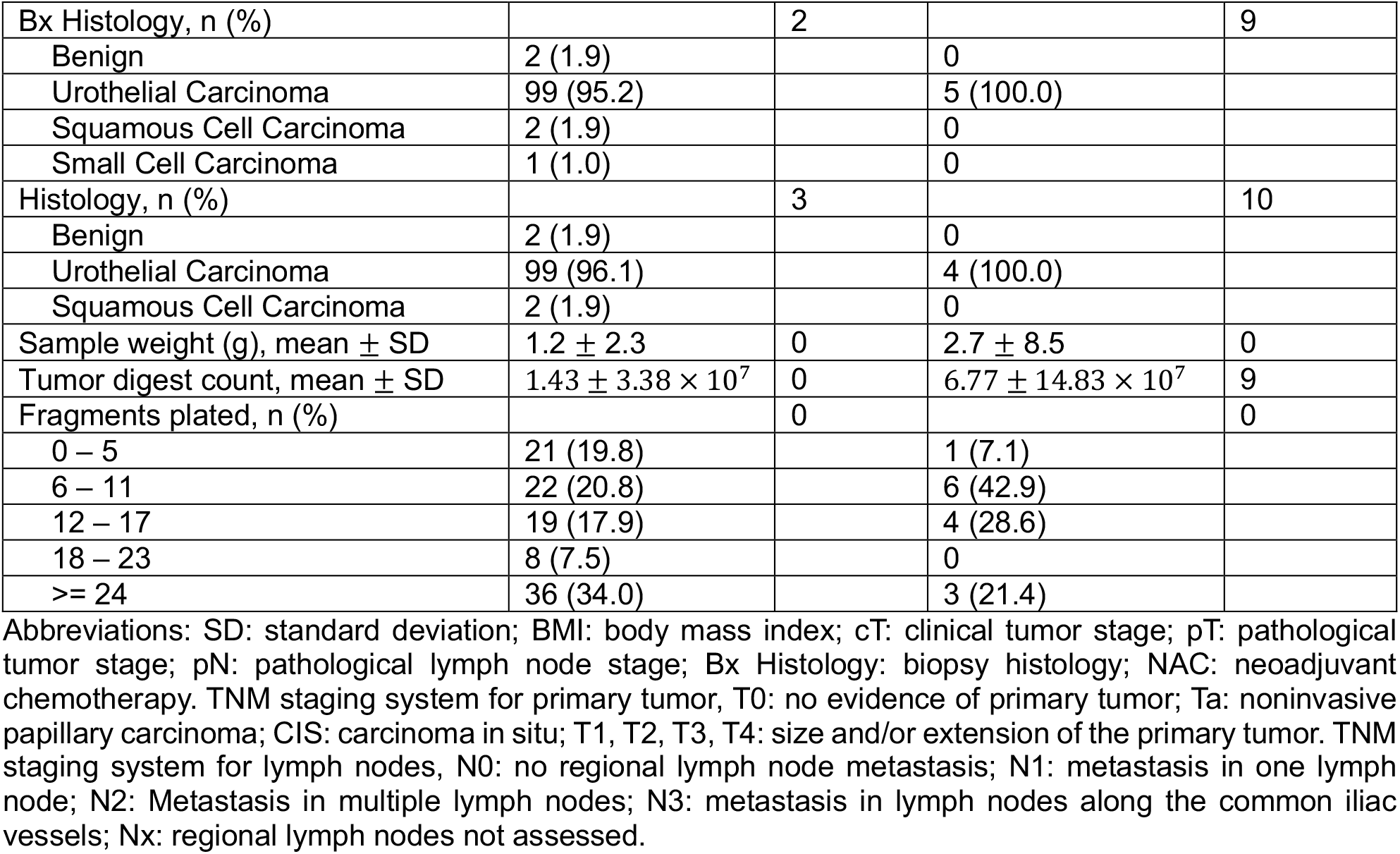
Description of Study Population.

The PETIL pipeline shown in Figure 1 consists of four steps: (i) data pre-processing that includes splitting data 70/30 into training and testing cohorts, data values normalization and imputation of the missing data, removal of features that are correlated, and analysis of ML classification methods for which the data sample size is adequate to drawn predictive conclusions; (ii) selection of a smaller subset of features that are robust in making predictions; (iii) model training that includes learning hyperparameters for the chosen ML predictor and determining the decision boundary threshold to account for imbalanced data; and (iv) generating predictions and calculating the performance metrics. After training, PETIL has established a minimal list of predictive features, a ML classifier adequate to the given data, and the decision threshold for the imbalanced data. Subsequently, PETIL was applied to stratify patients in the testing cohort. Finally, the performance metrics were computed, including accuracy, sensitivity, and specificity.

**Figure 1.**
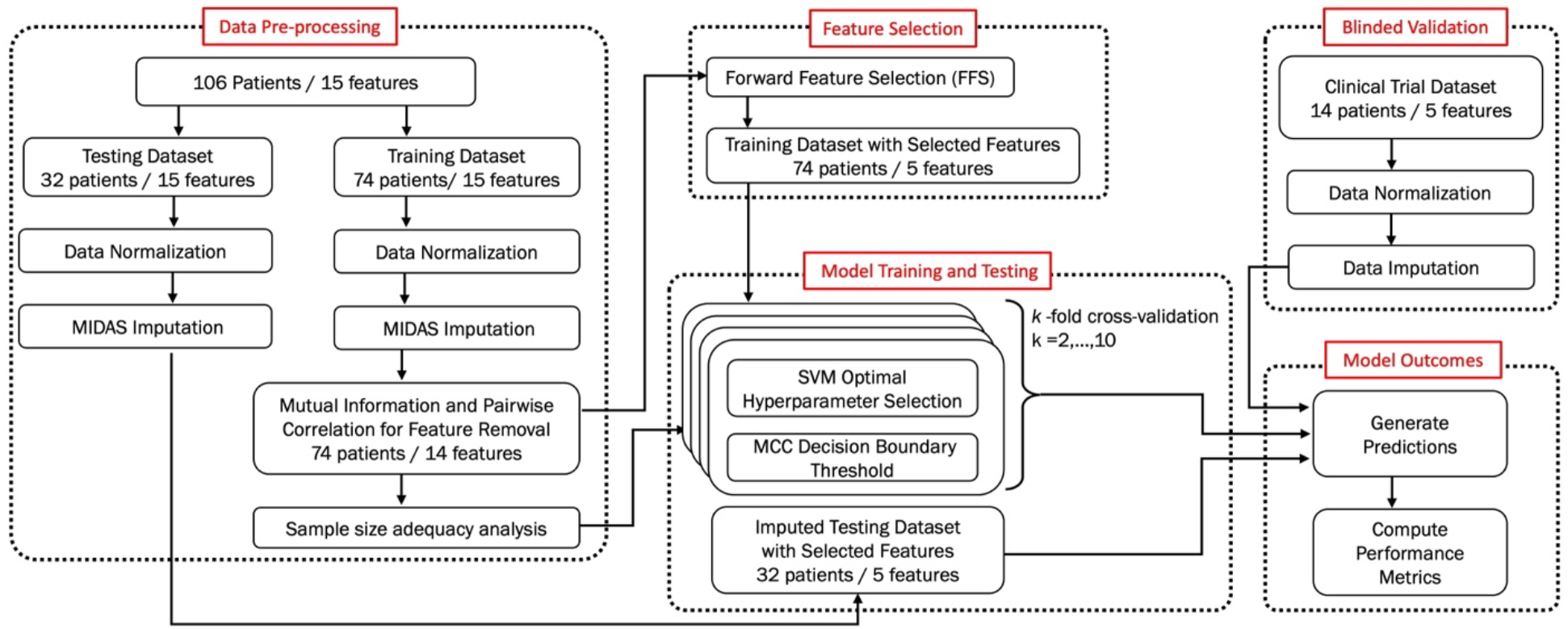
The PETIL pipeline. Abbreviation: PETIL: Predictor of the Expansion of Tumor Infiltrating Lymphocytes; MIDAS: Multiple Imputation with Denoising Autoencoders; FFS: Forward Feature Selection; SVM: Support Vector Machines.

For additional validation, the same type of data was collected from the phase I clinical trial “Intravesical Adoptive Cell Therapy w/ TIL for BCG Exposed High Grade NMIBC”, that was conducted at Moffitt Cancer Center between 2023 and 2025 (NCT05768347, (13)) under the protocol approved by the Advarra Institutional Review Board (MCC21894). Informed consent was also obtained from all patients prior to data collection. The summary of this data is also shown in Table 1. However, the corresponding TIL growth status was blinded until after the PETIL predictor was developed and the predictions for this cohort were generated. This dataset was also subject to data normalization and imputation of the missing data before PETIL generated predictions. Even if the clinical trial dataset was small, it was used as a blinded validation cohort due to the lack of a validation dataset from another institution, as the dataset we used here is the biggest collection of bladder cancer TIL specimens in one institution.

### Data normalization and imputation

After splitting 70/30 the retrospective dataset into training (74 patients) and testing (32 patients) cohorts, both were normalized separately using *MaxAbsScaler*, a class in the Scikit-learn Python library (17). The total proportion of missingness in the overall dataset was 4.4%, with 7 features having between 2 and 33 missing entries (Table 1). To address the issue of missing entries, we employed the Multiple Imputation with Denoising Autoencoders (MIDAS (18)) method, a deep learning approach that can generate imputed values while preserving interrelations among all features. MIDAS was applied to the training and testing cohorts separately and learned and imputed admissible values for the missing data. For the clinical cohort used for blinded validation, the data was normalized and all missing entries were imputed with zero value due to the small size (n = 14) of this cohort.

### Identification of pairwise collinear features

Using the Pearson’s pairwise correlation coefficient and setting the collinear threshold to 0.6 or higher (high positive correlation) and −0.6 or lower (high negative correlation), we tested all 15 features in the imputed training dataset to identify collinear features (Figure 2A). Next, we ranked the 15 features according to their Mutual Information (MI (19)) scores (Figure 2B). The Feature ‘cT or pT’ was the only one that met the collinearity threshold with either ‘pT’, ‘cT’, or ‘Surgery’ features, but due to lower MI score for ‘cT or pT’, this feature was removed from further consideration (Figure 2C). This reduced the dimension of the feature space from 15 to 14.

**Figure 2.**
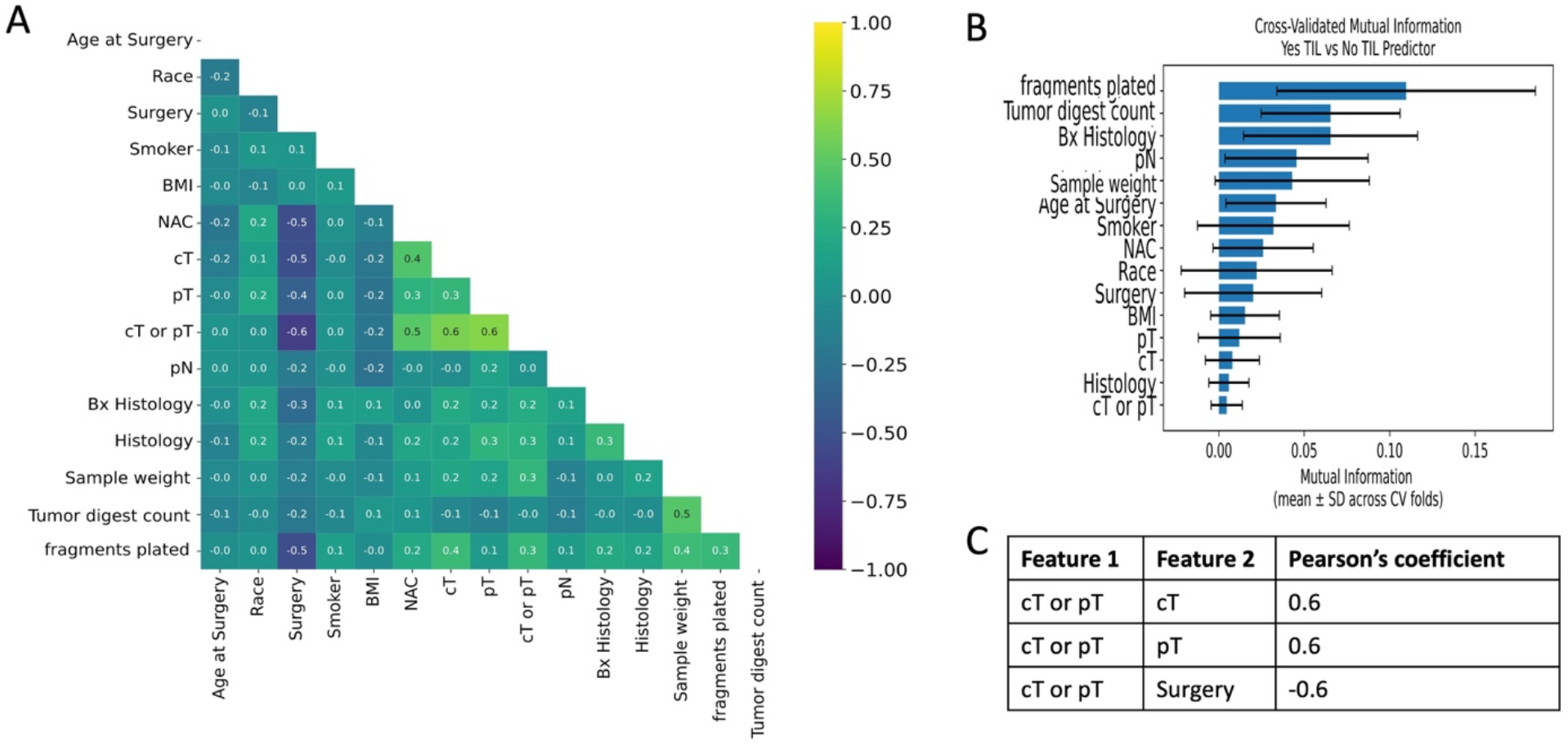
Identification of collinear features. **A.** Pearson’s Pairwise correlation for all 15 features. **B**. Mutual Information score for all 15 features **C**. The collinear features from **A** that met the cut-off threshold, due a lower MI score in **B** feature 1: ‘cT or pT’ will be removed.

### Sample size adequacy analysis

We conducted a sample size adequacy assessment by using the learning curves analysis (20, 21) to identify an appropriate ML classifier for the given task and given data sample. We showed that for the training dataset of size 74, the support vector machines (22–24) with the radial basis function kernel (RBF-SVM) were either optimal or performed better than the logistic regression (25), gradient boosting (26), and random forest (27) classifiers. The learning curves were obtained by stratified k-fold cross-validation on subsets of the training dataset containing from 10% to 100% of the available training dataset and evaluating performance on the held-out validation dataset using the RBF-SVM classifier (Figure 3). The learning curves show high validation scores for most of the sampled subsets of the training dataset. The low variance between the training scores and the validation scores, as well as the near convergence in the validation scores, indicates high generalization to new data and adequacy of the 74-training dataset for classification task. The learning curve analysis for the remaining three classifiers is shown in S1 Figure.

**Figure 3.**
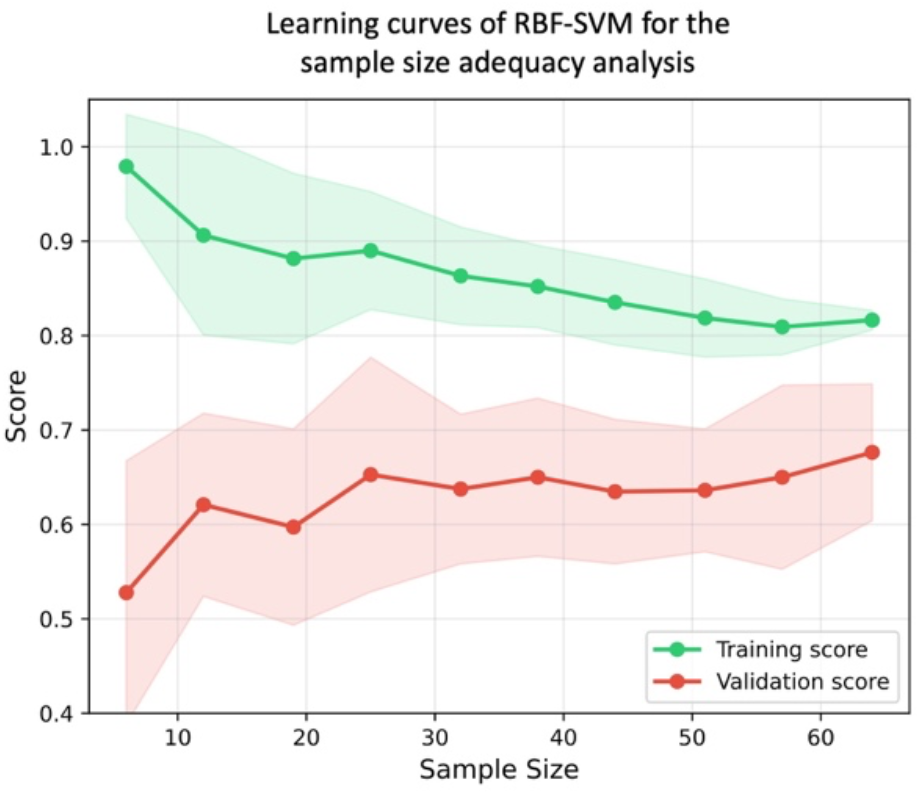
Sample size adequacy analysis for the RBF-SVM predictor. Learning curves show training (green) and validation (red) performance across training datasets of varying sample sizes. Shaded regions represent standard deviation across 8-fold cross-validation. The validation curve low variance with the training curve indicates that the classifier does not overfit to the 74-training set with 14 features. The validation curve show that performance analysis could be improved with more training data.

### Identification of a combination of robust predictive features

The 14 features commonly collected in the clinic were used as a base for identifying a smaller set of robust predictive features. The base set included 4 demographic, 7 clinical, and 3 biological tumor specimen-based features, all listed in the Materials and Methods section. Using the training dataset, we implemented the forward feature selection (FFS) method (28) to identify a subset of features that can distinguish between the *Yes-TIL* and *No-TIL* classes. FFS determined a pre-ranking of the 14 features by calculating the feature importance score (FIS) (27, 29, 30) using the random forest (RF) classifier on a stratified cross-validated 10 subsamplings of the training dataset (Figure 4A). Next, we used each fold of the 10 stratified cross-validated subsamples of the training dataset to train the RF classifier, where we sequentially added each feature in the order of its ranking and validated the addition of each feature on the holdout in each fold. This process of adding features one at a time yields a non-strictly increasing curve of the RF accuracy on the holdout sets. There may be local regions of downward fluctuations due to noisy features. To mitigate these fluctuations, the validation accuracy curve was converted to a monotonically increasing curve, retaining the maximal accuracy observed up to each feature addition step. The optimal number of features was determined using a stopping criterion that checks for plateau in the validation accuracy curve (31), beyond which additional features provide minimal incremental gain in accuracy (Figure 4B).

**Figure 4.**
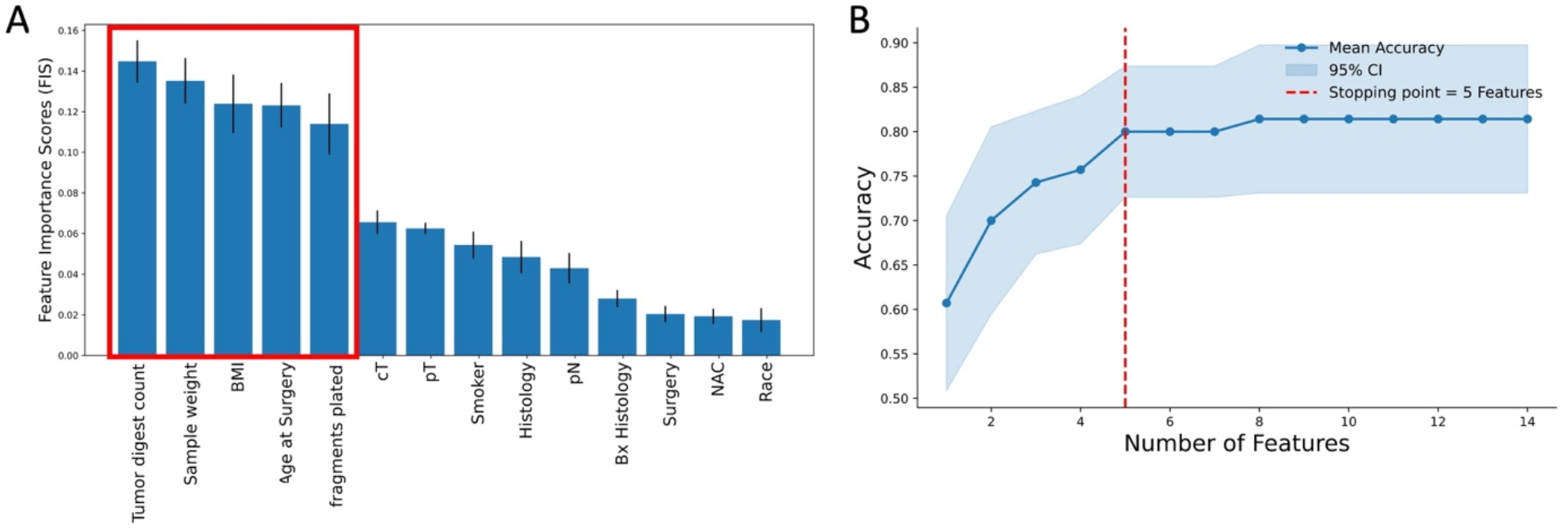
Feature selection. **A.** The 14 features were pre-ranked using the FIS values obtained from training the RF algorithm on a stratified cross-validated subsamples of the training dataset. **B**. The FFS algorithm calculated the validation accuracy of sequentially updated features with high FIS rank and then using a stopping criterion, it identified 5 robust predictive features.

The FFS algorithm determined 5 robust predictive features: (i) two demographic features (age at surgery, BMI) and (ii) three biological tumor specimen-based features (number of fragments plated, sample weight tumor, tumor digest count) shown boxed in Figure 4A. The distributions of the robust predictive feature values in the training and testing cohorts are shown in S2 Figure.

### Learning of the predictive SVM and MCC hyperparameters

The PETIL classifier of choice is the RBF-SVM because the learning curve analysis indicated its superior data generalization and adequacy of the size of the training dataset for classification in comparison with other classifier, such as logistic regression, gradient boosting, and random forest. Another advantage of the RBF-SVM classifier is its ability to capture interactions between predictive features that are potentially multidimensional and nonlinear (23, 24). Using the training dataset with the selected 5 robust predictive features, we learned the optimal classification hyperparameters using a *k*-fold cross-validation (for *k* = 2, …,10). For each *k*, we used the Matthews correlation coefficient (MCC) to determine the best decision boundary threshold so as to account for the imbalance in the dataset. For each *k*, we identified the best RBF-SVM hyperparameters (*C*, γ). Here, *C* specifies the width of the margins for avoiding data misclassification, while γ controls the nonlinearity of the decision boundary hyperplane. The best *C* and γ were determined by (i) minimizing the difference between the cross-validation training accuracy and holdout accuracy; (ii) maximizing the cross-validation training accuracy; and (iii) minimizing the value of *C*. We determined the optimal *k=5*, and the corresponding optimal hyperparameters values were *C* = 2.520 and γ = 0.02 (S3 Figure). The optimal MCC decision boundary threshold was 0.629.

### Evaluating the predictive performance of PETIL

The developed optimal RBF-SVM-MCC model was first verified using the testing dataset and was subsequently validated using the blinded validation (clinical trial dataset). The testing cohort comprised 32 patients, and the PETIL predictions yielded the accuracy of 0.750, the true positive rate (sensitivity) of 0.762, and the true negative rate (specificity) of 0.727 (Figure 5A). The area under the curve (AUC) for the testing cohort was 0.740 (Figure 5B). The clinical trial dataset comprised 14 patients and all successfully had TIL expansion. PETIL predicted this outcome with an accuracy of 0.857 (12 out of 14 correct predictions).

**Figure 5.**
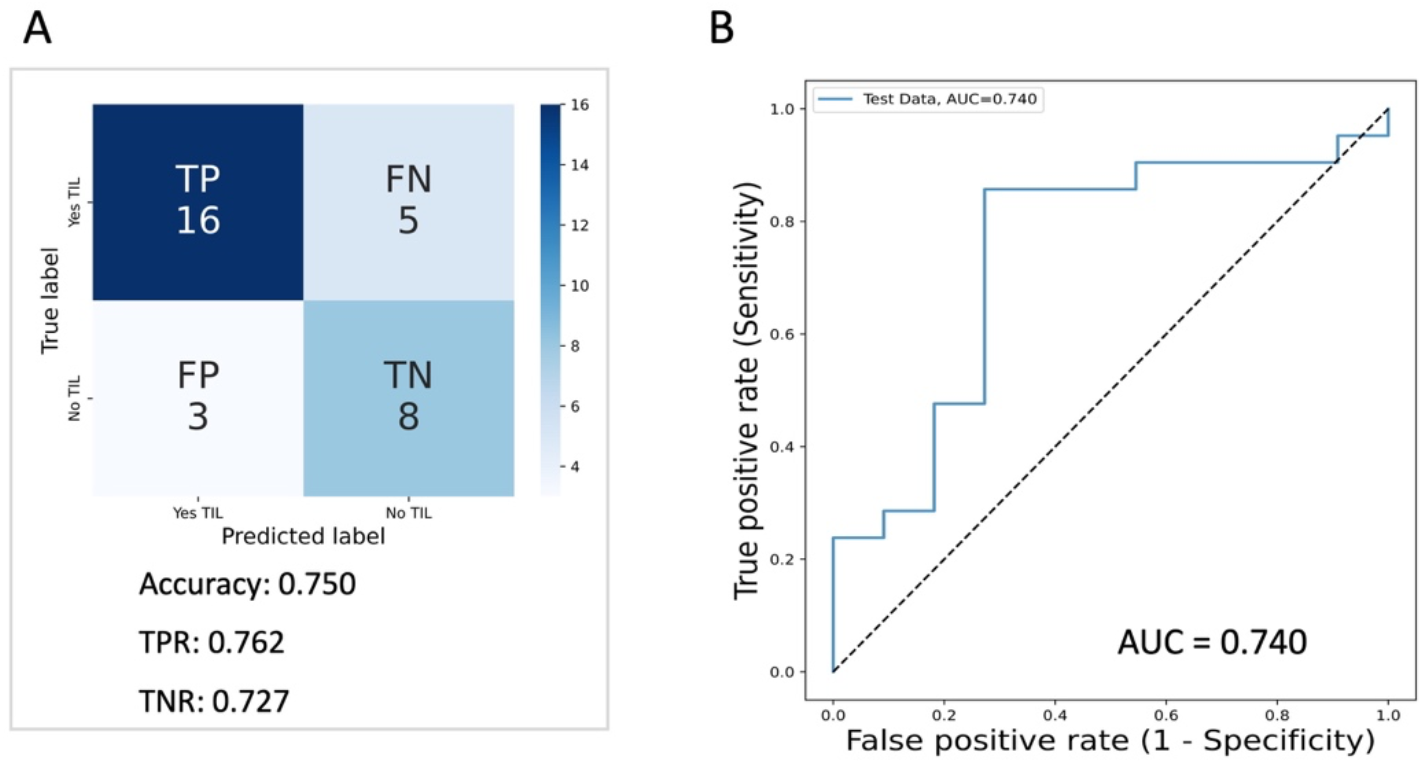
Model performance analysis. **A**. The confusion matrix of PETIL predictions for the testing dataset with the indicated performance metrics: the accuracy, true positive rate (TPR), and true negative rate (TNR). **B**. The receiver-operating characteristic (ROC) curve generated for PETIL predictions for the testing dataset with the indicated area under the curve (AUC). The dashed line represents a random classifier. Abbreviations: PETIL, Predictor of the Expansion of Tumor Infiltrating Lymphocytes.

## Discussion

Despite recent advancements in treatment of non-muscle invasive bladder cancer (32), especially once patients fail BCG (Bacillus Calmette-Guerin) a standard intravesical immunotherapy, long-term results are modest, with low durable treatment response rates and cystectomy-free rates far from ideal (33, 34). These modest results show a significant unmet need for further development of therapies for bladder cancer patients, preferably treatments that can be delivered locally with minimal associated adverse events and that can provide durable responses. Because bladder tumors have a high mutational burden, they are good candidates for adoptive cell therapy with autologous tumor-infiltrating lymphocytes (ACT-TIL), in which TILs are expanded and activated *ex vivo*, and then reinfused into the cancer patient as a personalized form of therapy. For bladder cancer patients, there is also a unique opportunity to deliver TILs locally, through intravesical administration, without systemic cytotoxic chemotherapy to induce lymphodepletion. Although ACT with TIL holds a great promise, we have previously shown that TIL expansion from the resected bladder tumors failed in about 30% of the patients (12). Moreover, *ex vivo* expansion of TIL takes about 4 weeks after tumor resection, thus early assessment of the outcome of TIL expansion could benefit the patient. Early identification of patients who could benefit from ACT-TIL is essential for optimizing treatment strategies and in preventing delays in next appropriate treatment. For example, if a patient is predicted to have a negative TIL expansion, other therapies could be considered. This improved selection of patients for ACT-TILs may prevent excessive treatment-related costs and delays in treatment of patients whose bladder cancer specimens would not yield successful TIL growth.

Here, we conducted a retrospective analysis of multimodal data of 106 bladder cancer patient specimens and developed the PETIL model for predicting whether a given tumor will be suitable for the successful expansion of TILs. One of the important features of this method is the capacity to first identify which data combinations are important for making predictions from a larger dataset in hand that has been already collected in clinic. Thus, our method does not require a predefined set of data features, but it learns which data features are robust in predicting the outcome from the available data. As a result, this method is optimized for the local data. Once the combination of robust predictive features was identified, the PETIL predictor showed favorable performance metrics on both the testing cohort that was a part of the retrospective database (AUC = 0.740) and the validation cohort that came from an active clinical trial (Accuracy of 0.857, correctly predicted 12 out of 14 TIL expansions).

Another important feature of the PETIL classifier is the capacity to make successful predictions using cohorts of a medium size (74 datasets for a training cohort and 32 for a testing cohort). This is in contrast to other ML methods, and deep learning methods in particular, that require large datasets for training. A limitation in our study is the lack of an external validation dataset from another institution. However, we utilized the phase I clinical trial dataset (NCT05768347) as a blinded validation cohort. To our knowledge, the combined dataset we used (11, 12) is the biggest collection of bladder cancer TIL specimens in one institution, and we are continuing to accrue bladder tumors from patients at Moffitt Cancer Center to include in future studies.

In conclusion, a novel ML model was developed based entirely on local data already collected in clinic without the need of acquiring additional data features. This model was able to predict TIL expansion with favorable performance. While further studies with larger cohorts of data are still desirable, PETIL predictor shows a great promise to be used as a clinical-supportive tool to help with stratifying patient eligibility for immune-based therapies, such as the ACT-TIL.

## Materials and Methods

### Clinical and experimental data

The prospective database of 106 adult patients was created between 2015 and 2022 by collecting clinicopathologic and specimen information of bladder cancer patients undergoing surgery at Moffitt Cancer Center. Data collection protocols and all experimental protocols were approved by the Advarra Institutional Review Board (MCC18142 and MCC20106) and performed in accordance to IRB guidelines. Informed consent was obtained from all patients prior to data collection. Additional data for 14 adult patients was collected from the phase I clinical trial (NCT05768347) conducted at Moffitt Cancer Center between 2023 and 2025 under the protocol approved by the Advarra Institutional Review Board (MCC21894). Informed consent was also obtained from all patients prior to data collection. Bladder tumors were included if they were larger than 1 cm and if there was tissue specimen available after pathological diagnosis. Tumors specimens were collected from transurethral resection of bladder tumor (TURBT) or radical cystectomy. Tumor infiltrating lymphocytes were expanded from resected surgical tumors as previously described (12). Tumors were minced into fragments, placed in individual wells of a 24-well plate, and propagated in media containing 6000 IU/mL IL-2 for up to four weeks. As cell confluence was reached, each well was split into additional wells. Expansion of TIL was considered successful if at least 2 wells were confluent at the end of four weeks. These cases were categorized as *Yes-TIL*. All other cases were categorized as *No-TIL*. This established database that was used for retrospective data analysis and ML-based predictions.

All datasets included information on 15 commonly collected features about patients’ (i) demographics: age at surgery, body-mass index (BMI), race, and smoker status; (ii) clinical characteristics: clinical tumor stage (cT), pathological tumor stage (pT), pathological lymph node stage (pN), type of surgery, prior neoadjuvant chemotherapy (NAC), type of radical cystectomy histology, type of cystoscopic biopsy histology, and cT/pT status; and (iii) specimen information: tumor sample weight, number of fragments plated for TIL expansion, and tumor digest count. In total, 56 patients were missing data in some considered features. We used the MIDAS imputation method, since it can handle both numerical and categorical data features. For every patient, information was recorded on whether the resected tumor sample caused positive TIL growth. For the clinical trial validation cohort, the TIL growth status was blinded until after the PETIL predictor was developed and the predictions for this cohort generated. The clinical trial dataset was used as a validation cohort due to the lack of a validation dataset from another institution, since the dataset we used here is the biggest collection of bladder cancer TIL specimens in one institution.

### Description of the ML classification problem

Our goal was to identify a predictive subset of features that differentiates between two targets in a binary classification task, and to provide metrics of success for such data stratification. Let the dataset **X** consists of Mdatapoints: **X**= [**X**_1_,**X**_2_,…,**X**_M_]^T^, where each data point has **p** features: = 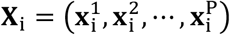 for i ∈ {1,2,…,M}, and for each i, the corresponding target is the binary class y_i_ ∈ {−1,1} The goal was to find the minimal subset of features of sizeQ, where Q<**p**, that divides the dataset **X** into distinct binary classes. We used a data-informed approach. First, we split **X**into training and testing cohorts. Using the training cohort, we implemented feature selection method to identify predictive features. Next, we determined an optimal machine learning classifier using the sample size adequacy methods. Subsequently, we found the optimal hyperparameters for the classifier using a cross-validation technique and also learned the optimal decision boundary threshold that corrects for imbalance in the binary classes in **X**. Finally, this classifier was applied to the testing cohort to assess the prediction metrics.

### Data normalization

Data normalization is performed to ensure that all features can contribute equally. In order to prevent data leakage between training and testing cohorts, we split the overall data before data normalization, and perform normalization on the training and testing cohorts separately. We used the *MaxAbsScaler*, a class in the scikit-learn Python library (17) which translates each feature independently to have a maximal absolute value of 1.0, preserving data distribution but only linearly scales down each feature. Data normalization was also used for the clinical trial cohort.

### Multiple Imputation framework for data retention

Multiple imputation is a statistical method to handle missing data. Let **X**∈ ℝ^M×p^ be a data matrix with observed entries **X**_obs_ and missing entries **X**_miss_. Under the assumption that data are missing at random (MAR) or completely at random (MCAR), the multiple imputation replaces all entries in **X**_miss_ with imputed values that preserve the interrelations in **X**_obs_. We use here the multiple imputation with denoising autoencoders (MIDAS) method (18), which is a scalable deep learning-based technique that employs a class of unsupervised neural networks known as denoising autoencoders (35) and Monte Carlo dropout to generate multiple imputation of the missing data with realistic uncertainty quantification. In our training dataset 32 out of 74 patients were missing data in at least one feature in the training dataset, and 14 out of 32 patients were missing data in at least one feature in the testing dataset. In either cohort, the MIDAS data imputation method was used to impute missing data. The imputed training dataset was used to identify robust predictive features and to apply the RBF-SVM and MCC algorithms.

### Mutual information measure for nonlinear dependencies

Mutual information (MI) is a non-parametric measure of statistical dependency between the dataset **X**∈ ℝ^M×p^ and the predicted binary class Y ∈ ℝ^M×1^, where Mis the number of patients and p is the number of features. MI captures nonlinear dependencies in high-dimensional data, which makes them robust for measuring feature relevance in discrete or categorical datasets (36, 37). A discrete MI is computed as follows: 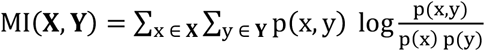 Where p(x,y) is the joint probability of **X** and Y, while P(x)and p(y)are the marginal probabilities of **X** and Y, respectively.

### Learning curve method for determining sample size adequacy

To assess training dataset adequacy for a classification task of distinguishing TIL expansion status, we performed learning curve analysis using stratified k-fold cross-validation, where we repeatedly subsampled the training data at different sample sizes. We trained four models (Logistics Regression, Random Forest, Gradient Boosting, and RBF-SVM) on subsets of the training dataset from 10% to 100% of the available training dataset and evaluated performance on the holdout validation dataset.

### Forward feature selection method for identification of predictive features

The feature selection was performed to identify the minimal predictive subset of features used in the data classifier. Using the Forward Feature Selection (FFS) method (28), we built a stratified cross-validated 10 subsamplings of the training dataset. On each fold, we trained a random forest (RF) classifier and obtained feature importance score (FIS) ranking for each of the features. Next, the features were added sequentially in the order of pre-ranking, and the predictive accuracy of the RF classifier was evaluated using the holdout validation set. To mitigate fluctuations due to noisy features, accuracy curves were converted to a monotonic increasing curve, retaining the maximum accuracy observed up to each feature addition step. The optimal number of features for each fold was determined using a stopping criterion which identifies the point beyond which additional features provide minimal incremental gain in accuracy. This stopping criterion also prevents overfitting to the training data. The mean accuracy curve and 95% confidence interval across all folds were used to visualize feature contribution to the predictiveness of the model and to determine final feature selection.

### Support vector machine model for learning the optimal classification rules

For each data point 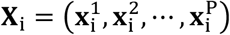 in **X**∈ ℝ^M×p^, a binary ML classifier learns a corresponding prediction y_i_ in Y ∈ ℝ^M×1^. The accuracy of this prediction depends on how well the classifier identifies an optimal decision boundary That separates the **X** into the distinct binary classes. However, the interactions between the predictive features are often multidimensional and nonlinear. A support vector machine (SVM) model with the nonlinear Radial Basis Function (RBF) kernel: k(**X**_i_,**X**_j_)=exp(−γ∥**X**_i_ –**X**_j∥_^2^)(23, 38) can potentially capture this complex interaction between features as it learns to distinguish between the distinct classes in the training dataset and generalize to new dataset. First, all data was split 70/30 into the training and testing cohorts. Then, the optimal RBF-SVM hyperparameters (C, γ) were identified by performing k-fold cross-validation (CV) on the training dataset, for k = 2, …,10. Here C specifies the width of the margins for avoiding data misclassification and γ controls the nonlinearity of the decision boundary hyperplane. For each k, the best C and γ were determined by (i) minimizing the difference between the cross-validation training accuracy and testing accuracy; (ii) maximizing the cross-validation training accuracy; and (iii) minimizing the value of C. The optimal values of C and γ were obtained by computing the Matthews correlation coefficient (MCC) for each cross-validation (k = 2, …,10) and then choosing C and γ for which MCC is maximal, and C is minimal.

### Matthews Correlation Coefficient algorithm for adjusting imbalanced datasets

The Matthews correlation coefficient (MCC) statistical test (39, 40) was used to determine the optimal decision boundary threshold (for *Yes-TIL* and *No-TIL* classifications) that accounts for imbalances in the training dataset. Usually, this threshold is set to 0.5, because it is assumed that SVM works with balanced data (i.e., similar numbers of data fall into each class). For the imbalanced dataset, this threshold has to be adjusted. This adjustment was done by testing thresholds between 0.01 and 0.99 by separating the training cohort data into *Yes-TIL* or *No-TIL* classes based on whether their RBF-SVM–generated prediction probabilities exceeded the given threshold. For each threshold, MCC was calculated on the labeled prediction probabilities. The threshold with the maximum MCC was called optimal and was used as the decision boundary threshold to generate predictions on the testing cohort. In this approach, the magnitude of the decision function **w**^T^**X**_i_ + b for each **X**_i_ were extended to probability estimates using the scikit-learn library (41), with option ‘probability=True’. A detailed algorithm is presented in S1 Algorithm.

### Performance metrics for PETIL evaluation

To assess performance of the classification protocol on the testing dataset, the following performance metrics were used for evaluation: (1) true positive rate (TPR) or sensitivity, is the percentage of correctly classified positive instances: TPR=TP/(TP+FN); (2) true negative rate (TNR) or specificity, is the percentage of correctly classified negative instances: TNR=TN/(FP+TN); (3) accuracy is the percentage of correctly classified positive and negative instances: accuracy=(TP+TN)/(TP+FN+FP+FN); (4) area under the receiver operating characteristics curve (AUC/ROC or AUC) measures the ability to discriminate between positive and negative cases and ranges from 0.5 (coin toss) to 1.0 (perfect classification), when ROC curve shows tradeoffs between TP and FP. Here, TP (true positive) is the correctly classified data, TN (true negative) is the correctly classified data, FP (false positive FP) is the misclassification of the positive class, and FN (false negative) is the misclassification of the negative class.

## Supporting Information Captions

**S1 Table:** Pearson Correlation Coefficient between each of 15 individual data features and the TIL growth status

**S1 Figure**: **Learning curve analysis for different ML classifiers. A**. Learning curves for the logistic regression (LR) model. Low validation scores indicate that LR is not an adequate classifier. **C**. Learning curves for the random forest (RF) method. High variance between the training and validation scores shows that RF is not an adequate classifier. **D**. Learning curves for the gradient boosting (GB) method. High variance between the training and validation scores shows that GB is not an adequate classifier. In each analysis, sample proportions ranging from 10% to 100% of the training dataset were sampled multiple times. A stratified k-fold cross-validation was used to obtain the training (shown in green) and validation (shown in red) performance curves across varying training dataset sample sizes. Shaded regions represent standard deviation across 8-fold cross-validation.

**S2 Figure: Distributions of robust predictive features**. Distributions of the *No-TIL* (blue) and *Yes-TIL* (orange) classes for the training (left) and testing (right) cohorts for five robust predictive features: **A**. Tumor sample weight, **B**. Tumor digest count, **C**. Age at surgery, **D**. BMI, **E**. Number of fragments plated.

**S3 Figure: Optimal hyperparameter search**. Hyperparameter space for parameters C and γ for the RBF-SVM with cross-validation k=5, for testing (left) and training (right) cohorts.

**S1 Algorithm:** Algorithm for determining the optimal decision threshold for radial basis-kernel function for the support vector machine (RBF-SVM) and the Matthews correlation coefficient (MCC) methods.

## Acknowledgments

Assistance in data collection and de-identification was provided by Ms. Suzanne McFarland, an honest broker at Moffitt Cancer Center.

## Ethics Statement

These studies were reviewed and approved by the Moffitt Scientific Review Committee (SRC) and Advarra Institutional Review Board (MCC18142, MCC20106, and MCC21894) and performed in accordance with IRB guidelines. Informed consent was obtained from all patients prior to data collection.

## Code availability

Computational code used in this study is available from the following GitHub depositories: https://github.com/okayode/Predictor_of_the_Expansion_of_TIL_project, https://github.com/rejniaklab/Predictor_of_the_Expansion_of_TIL_project

## Data availability

All data used in this study is available from the following GitHub links: https://github.com/okayode/Predictor_of_the_Expansion_of_TIL_project, https://github.com/rejniaklab/Predictor_of_the_Expansion_of_TIL_project

## Author Contributions

K.D.O., A.M.A., I.C., S.P.T., M.A.P., and K.A.R. participated in study design. A.M.A., S.B., B.B., S.P.T., and M.A.P. were responsible for data collection. K.D.O., A.K., S.P.T., M.A.P., and K.A.R. developed the methodology. K.D.O. and K.A.R. performed the formal analysis and wrote the initial draft of the manuscript, which was critically reviewed by A.M.A, A.K., S.P.T., and M.A.P. All authors approved the final version of the manuscript.

