## Supplemental Material for "PETIL: Predicting Expansion of Tumor Infiltrating Lymphocytes for the Adoptive Cell Immunotherapy in Bladder Cancers"

### Supporting Information

**S1 Table:** Pearson Correlation Coefficient between each of 15 individual data features and the TIL growth status

| Features | Pearson Correlation |
|---|---|
| Age at Surgery | -0.0892040899258394 |
| BMI | 0.04329824343984757 |
| Race | -0.16219222183503318 |
| Smoker | 0.06419727740004512 |
| cT | 0.22677980365766873 |
| pT | 0.05411552433036292 |
| pN | -0.023836393366219028 |
| Surgery | -0.21884238755804317 |
| NAC | -0.018377261547332296 |
| Histology | 0.11429218135123825 |
| Bx Histology | 0.21949168078954456 |
| cT or pT | 0.21908072935486692 |
| Tumor sample weight | 0.09418244178887078 |
| Number of fragments plated | 0.30320242877943865 |
| Primary tumor digest count | 0.13855431863926465 |

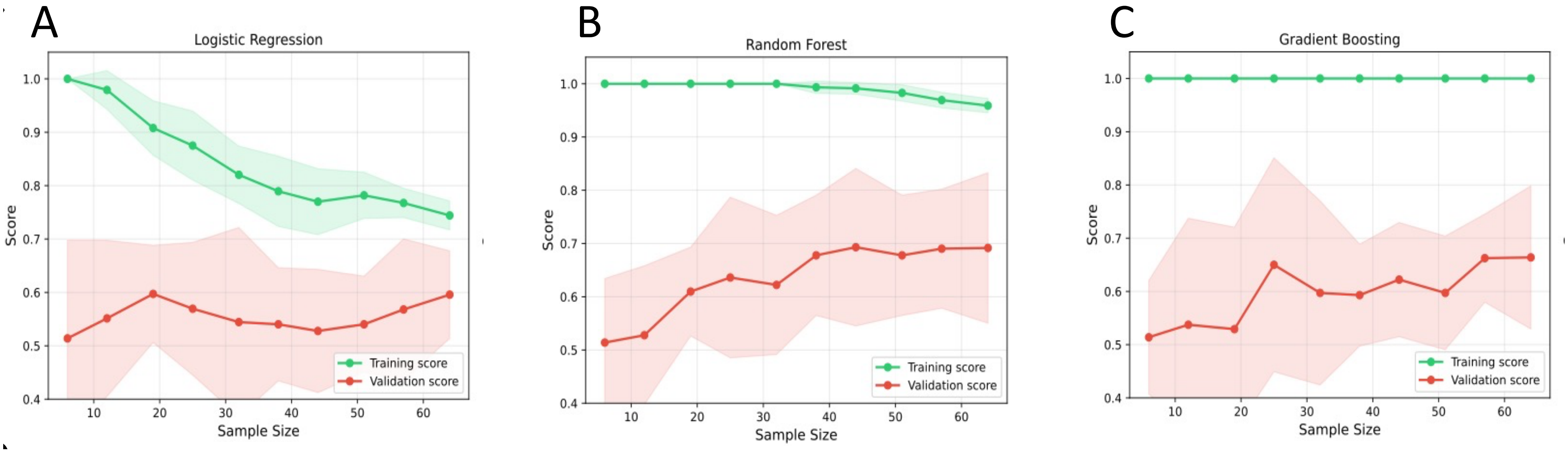


**S1 Figure**: **Learning curve analysis for different ML classifiers. A.** Learning curves for the logistic regression (LR) model. Low validation scores indicate that LR is not an adequate classifier. **C.** Learning curves for the random forest (RF) method. High variance between the training and validation scores shows that RF is not an adequate classifier. **D.** Learning curves for the gradient boosting (GB) method. High variance between the training and validation scores shows that GB is not an adequate classifier. In each analysis, sample proportions ranging from 10% to 100% of the training dataset were sampled multiple times. A stratified k-fold cross-validation was used to obtain the training (shown in green) and validation (shown in red) performance curves across varying training dataset sample sizes. Shaded regions represent standard deviation across 8-fold cross-validation.

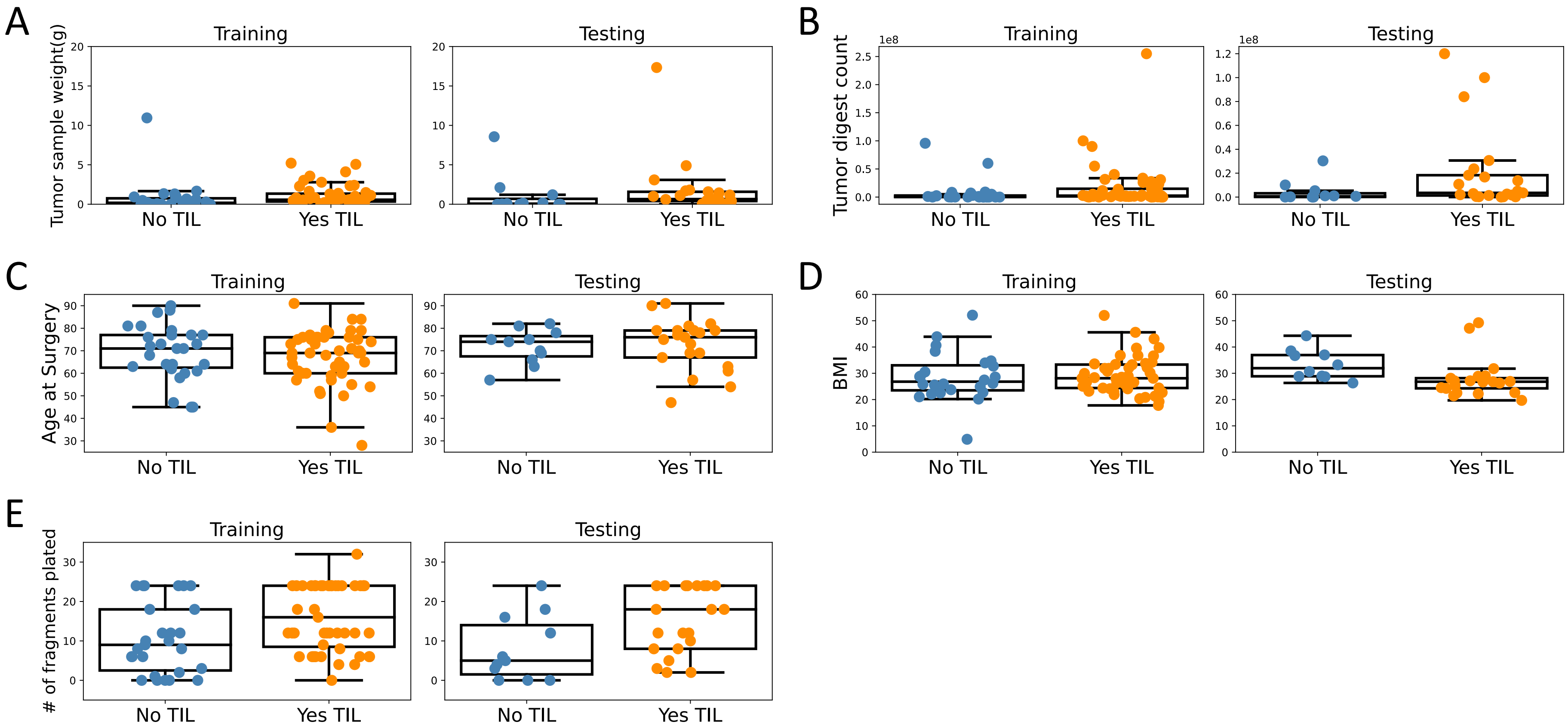


**S2 Figure: Distributions of robust predictive features.** Distributions of the *No-TIL* (blue) and *Yes-TIL* (orange) classes for the training (left) and testing (right) cohorts for five robust predictive features: **A.** Tumor sample weight, **B.** Tumor digest count, **C.** Age at surgery, **D.** BMI, **E.** Number of fragments plated.

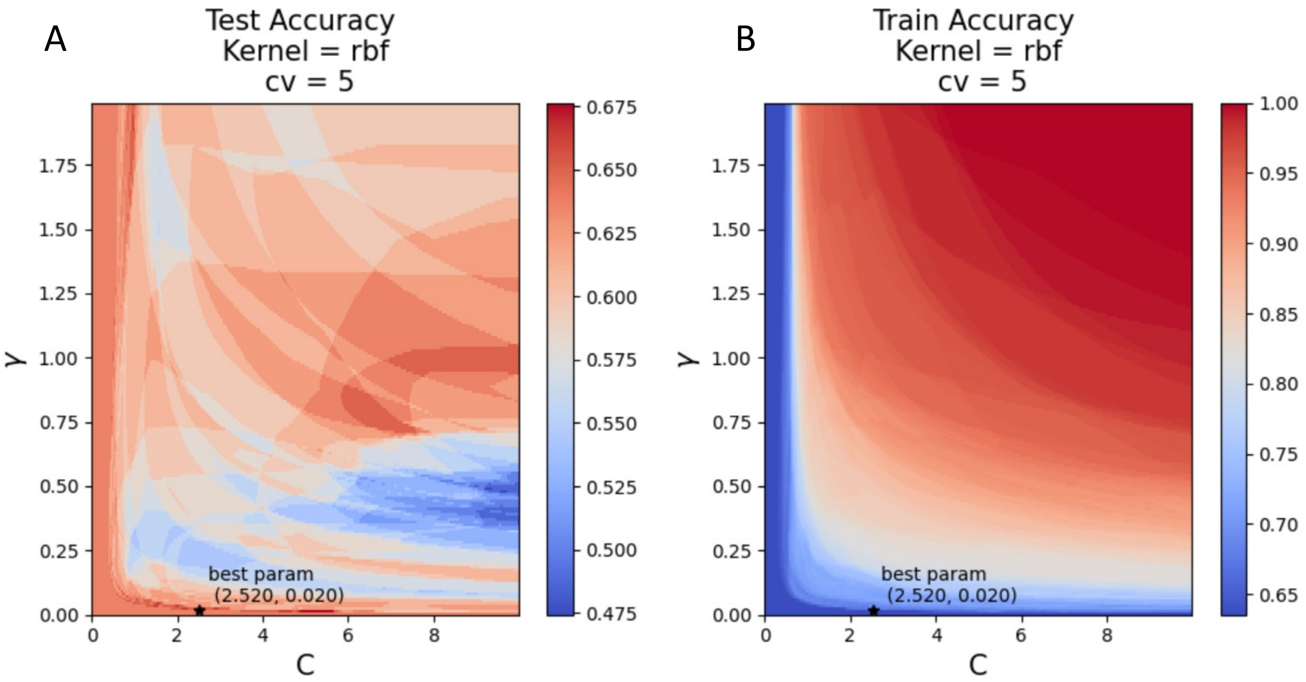


**S3 Figure: Optimal hyperparameter search.** Hyperparameter space for parameters C and γ for the RBF-SVM with cross-validation k=5, for testing (left) and training (right) cohorts.

**Algorithm 1:** Determine Optimal Threshold using RBF-SVM and MCC

**Input:** Labeled training data $\mathcal{D}_{\text{train}} = \{(\mathbf{x}_i, y_i)\}_{i=1}^{M}$, where $\mathbf{x}_i = \left(x_i^1, x_i^2, \ldots, x_i^P\right)$, $y_i \in \{-1, 1\}$

**Output:** Decision threshold $\theta^*$ that maximizes classification performance

Train a radial basis function (RBF) kernel SVM classifier $\mathcal{M}$ on $\mathcal{D}_{\text{train}}$ with optimal hyperparameters

For each input $\mathbf{x}_i$, compute the predicted class probability $\hat{p}_i = \mathbb{P}_{\mathcal{M}}(y_i = 1 \mid \mathbf{x}_i)$

**for** *threshold* $\theta$ *from* 0.01 *to* 0.99 *in steps of* 0.001 **do**

  Convert probabilities to binary predictions using:

$$\hat{y}_i^{(\theta)} = \begin{cases} 1 & \text{if } \hat{p}_i \geq \theta \quad \text{(`Yes TIL')} \\ -1 & \text{otherwise} \quad \text{(`No TIL')} \end{cases}$$

  Compute the Matthews correlation coefficient (MCC) between the predicted labels $\hat{y}^{(\theta)}$ and the true labels $y$

Identify the threshold that yields the best performance:

$\theta^* \leftarrow \arg\max_{\theta} \text{MCC}(\hat{y}^{(\theta)}, y)$

Apply $\theta^*$ to classify instances in the test set using probability outputs from $\mathcal{M}$

**S1 Algorithm:** Algorithm for determining the optimal decision threshold for radial basis-kernel function for the support vector machine (RBF-SVM) and the Matthews correlation coefficient (MCC) methods.
